# Apoptotic Find-me Signals are an Essential Driver of Stem Cell Conversion To The Cardiac Lineage

**DOI:** 10.1101/2021.06.21.449262

**Authors:** Loic Fort, Vivian Gama, Ian G. Macara

## Abstract

Pluripotent stem cells can be driven by manipulation of Wnt signaling through a series of states similar to those that occur during early embryonic development, transitioning from an epithelial phenotype into the cardiogenic mesoderm lineage and ultimately into functional cardiomyocytes ^1–4^. Strikingly, we observed that induced pluripotent stem cells (iPSCs) and embryonic stem cells (ESCs) undergo widespread apoptosis upon Wnt activation, followed by a synchronous epithelial-mesenchymal transition (EMT). The EMT requires induction of transcription factors *SNAI1/SNAI2* downstream of *MESP1* expression, and double knock-out of *SNAI1/2*, or loss of *MESP1* in iPSCs blocks EMT and prevents cardiac differentiation. Remarkably, blockade of early apoptosis chemically or by ablation of pro-apoptotic genes also completely prevents the EMT, suppressing even the earliest events in mesoderm conversion, including *EOMES, TBX6*, and *MESP1* induction. Conditioned medium from WNT-activated WT iPSCs overcomes the block to EMT by cells incapable of apoptosis (Apop-), suggesting the involvement of soluble factors from apoptotic cells in mesoderm conversion. Treatment with a purinergic P2Y receptor inhibitor or addition of apyrase demonstrated a requirement for nucleotide triphosphate signaling. ATP was sufficient to induce a partial EMT in Apop- cells treated with WNT activator. We conclude that nucleotides, in addition to acting as chemo-attractants for clearance of apoptotic cells can, unexpectedly, function as essential paracrine signals in mesoderm specification.

## Main

A mesenchymal-epithelial transition (MET) is an early and essential event during reprogramming by the Yamanaka factors *SOX2, OCT4, c-MYC* and *KLF4* ^5–7^. During reprogramming of mouse fibroblasts, SOX2 and OCT4 suppress *SNAI1/2* and *ZEB1/2*, which are key transcription factors that promote the reverse process, an epithelial-mesenchymal transition (EMT), while c-MYC represses TGFβ signaling and KLF4 induces epithelial gene expression ^8,9^. EMT occurs during gastrulation as epithelial epiblast cells ingress through the primitive streak^10^. EMT is obligatory for ESCs to differentiate into definitive endoderm^11^, while a sequential EMT/MET occurs during the conversion of the hepatocyte lineage^12^. However, the reason why these processes are required for reprogramming and differentiation remains unknown.

### Initiation of cardiomyocyte specification triggers apoptosis followed by an EMT

Human iPSCs (GM25256) used in this study expressed appropriate pluripotency markers (NANOG, OCT4, SOX2) and epithelial polarity proteins including SCRB, LLGL2, PAR-3, PKC-zeta and PALS1 (Supplementary Fig S1A). They assemble ZO-1 positive tight junctions (TJ) and E-cadherin positive adherens junctions (AJ), localize polarity proteins appropriately, and form cell monolayers with a cobblestone appearance (Fig 1A). Treatment with a WNT activator (CHIR 99021) for 48 hrs followed by treatment with a WNT signaling inhibitor (Fig 1B) was used to drive cardiomyocyte specification^2^. This protocol promoted exit from pluripotency and the sequential induction of mid-primitive streak, marked by induction of *EOMES* and *TBXT* (Bra/T) from 9 - 36 hrs; lateral mesoderm, marked by induction of *TBX6* and *MESP1* from 36 – 48 hrs, and cardiac mesoderm, marked by induction of *ISL1, NKX2.5* and *ANP*, for 48 – 72 hrs (Supplementary Fig S1B, C). Consistent with previous data^1,2^, spontaneous beating of immature cardiomyocytes was observed after 10-12 days (Supplementary Video 1). Cardiac specification was accompanied by loss of E-cadherin, and expression of Slug and Vimentin (Supplementary Fig S1D-E).

**Figure 1.**
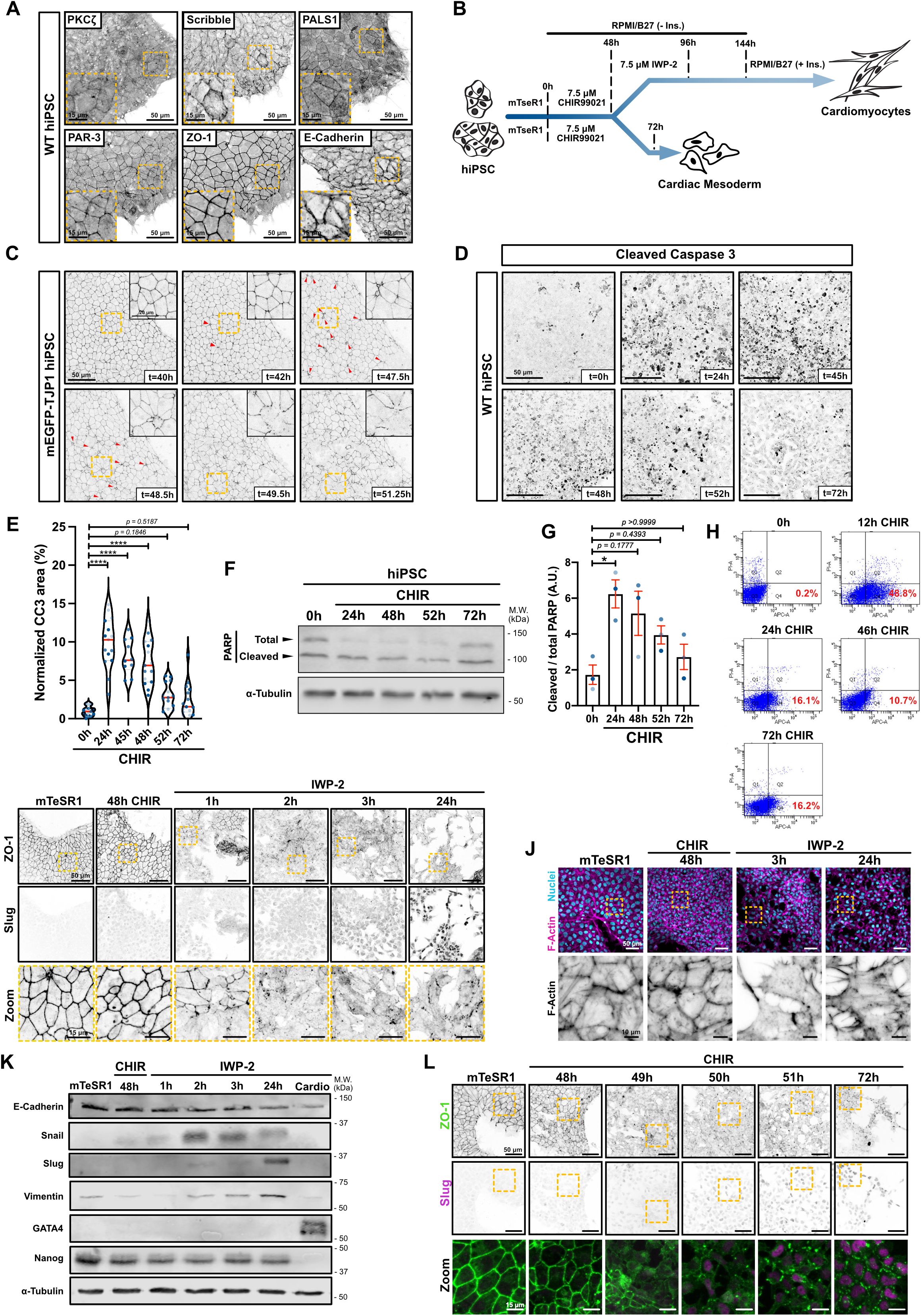
A) Immunofluorescence of hiPSC GM25256 stained for epithelial/polarity markers (Tight junctions: ZO-1 ; Adherent junctions: E-Cadherin ; Baso-lateral marker: Scribble ; Apical PAR complex: PKCζ, PAR-3 ; Apical Crumbs complex: PALS1). Maximum intensity projections are shown. Scale bar = 50 μm. Inset represents a magnified area (yellow dotted square). Scale bar = 15 μm. B) Schematic of GiWi differentiation protocol (top path) (conversion of hiPSC into cardiomyocytes); or alternative cardiac mesoderm with a prolonged incubation in CHIR-99021 (bottom path). C) Stills from Supp. Movie 1. mEGFP-TJP1 knock-in hiPSCs were imaged from 40h after CHIR treatment. Red arrows represent extruding cells. Scale bar = 50 μm. Magnified area (yellow dotted square) is shown on the right-hand corner. Scale bar = 20 μm. D-E) Representative immunofluorescence images of wildtype hiPSCs fixed at the indicated times after CHIR addition and stained for cleaved Caspase-3. Maximum intensity projections are shown. Scale bar = 50 μm (D). Violin plots quantifying the area of cleaved caspase 3-positive cells normalized to the cellular area. Independent biological repeats are color-coded (n=3, 5 random fields of view/repeat). (Median: plain red line – Quartiles: black dotted lines). Tukey’s multiple comparison was applied (**** P ≤ 0.0001) (E). F-G) Immunoblot of PARP cleavage in hiPSCs during CHIR treatment. Molecular weights (M.W.) are indicated in kDa (F). PARP cleavage was quantified by densitometry across 3 independent biological repeats (color-coded). Dunn’s multiple comparison was applied (* p < 0.05) (G). H) Cytometry gates from an Annexin-APC assay. Cells were treated with CHIR for different times, collected, stained and analyzed for Annexin (x-axis) and PI (y-axis). Percentage of Annexin-positive/PI-negative (Gate Q4 – Early apoptosis) is reported for each condition. I) Representative immunofluorescence images of hiPSCs (mTeSR1) or cells undergoing differentiation using CHIR/IWP-2 protocol, and stained for ZO-1 and Slug. Maximum intensity projections are shown. Scale bar = 50 μm. Magnified areas of the ZO-1 staining (yellow dotted square) are shown in the bottom row. Scale bar = 15 μm. J) Representative immunofluorescence images of wildtype hiPSCs fixed at the indicated times along the differentiation protocol and stained for F-actin (Phalloidin – Magenta) and nuclei (Dapi). Maximum intensity projections are shown. Scale bar = 50 μm. Magnified area of the F-actin channel (yellow dotted square) is shown. Scale bar = 10 μm. K) Immunoblots for EMT markers (E-Cadherin, Snail, Slug, Vimentin) during hiPSC differentiation to early cardiac mesoderm. Membranes were also blotted for GATA4 (cardiac specific marker) and Nanog (pluripotency marker). Molecular weights (M.W.) are indicated in kDa. L) Representative immunofluorescence images of hiPSCs (mTeSR1) or undergoing differentiation using prolonged CHIR incubation, and stained for ZO-1 and Slug. Maximum intensity projections are shown. Scale bar = 50 μm. Magnified area (yellow dotted square) is shown as a merge. Scale bar = 15 μm.

Strikingly, early induction of cardiomyocyte specification resulted in extensive cell extrusion (Fig 1C and Supplementary Video 2), caused by apoptosis that began within 12 hours, and slowly diminished over 2 days, as measured by cleaved caspase 3, cleaved PARP, and Annexin exposure (Fig 1 D-H). This was followed at 49-50 hrs by an abrupt wave of intercellular junction disassembly that occurred throughout the culture and was complete within ~2 hrs (Fig 1I; Supplementary video 2). Cell-cell contacts and cortical actin were lost, with acquisition of stress fibers and a spindle-shaped morphology (Fig 1 I,J). Surprisingly, expression of the AJ-marker E-cadherin did not diminish until ~24 hrs after junction disassembly (Fig 1K). Nonetheless, other markers of EMT were detected, including Snail and Slug, and the mesenchymal marker vimentin (Fig 1K, Supplementary Fig S1 D-E). Importantly, both apoptosis and EMT also occurred in hESCs that were driven along the cardiac mesoderm lineage, with increased PARP cleavage over the initial 2 days (Supplementary Fig S1F-G), followed by increased Snail and Slug (Supplementary Fig S1H). Importantly, the extensive apoptosis we observed is specific to Wnt activation, as treatment with retinoic acid does not stimulate cell death^13^.

The global EMT occurs almost immediately after addition of IWP-2 inhibitor in the standard protocol for cardiomyocyte induction (Fig 1I-K). However, the causal relationship between these two events was unclear. Therefore, we initiated mesoderm induction by addition of CHIR but withheld inhibitor at 48 hrs and did not remove the CHIR. Notably, the EMT still occurred on schedule in these cells (Fig 1L and Supplementary Fig S1I), demonstrating that it is independent of WNT inhibition and that, with astonishing fidelity, the iPSCs can synchronously time the EMT to a 1 – 2 hr window 2 days after WNT activation.

### EMT is driven by the induction of SNAI1 and SNAI2, downstream of MESP1

To investigate the mechanism of EMT, we first used RT-qPCR to analyze *SNAI1/2* induction and found that *SNAI1* peaks at ~50hrs while *SNAI2* peaks at about 72 hrs (Fig 2A), as also detected by immunoblot (Fig. 1K). We next asked if the observed EMT is dependent on *SNAI1* and/or *SNAI2*, by CRISPR/Cas9-mediated gene editing in the iPSCs (Fig 2 B,C). Knockout of either gene alone had little effect because of compensatory induction of the remaining gene (data not shown), but a double knockout (DKO) of *SNAI1/2* efficiently blocked the EMT (Fig 2D). Moreover, these cells did not continue down the cardiac mesoderm lineage. They did not express key cardiac markers such as *NKX2.5* and c*TNT* and showed a reduced expression of *GATA4* (Fig. 2E). We conclude that the EMT induced in response to WNT activation in iPSCs is driven by expression of Snail and Slug, and that these factors are essential for specification of the cardiac lineage.

**Figure 2.**
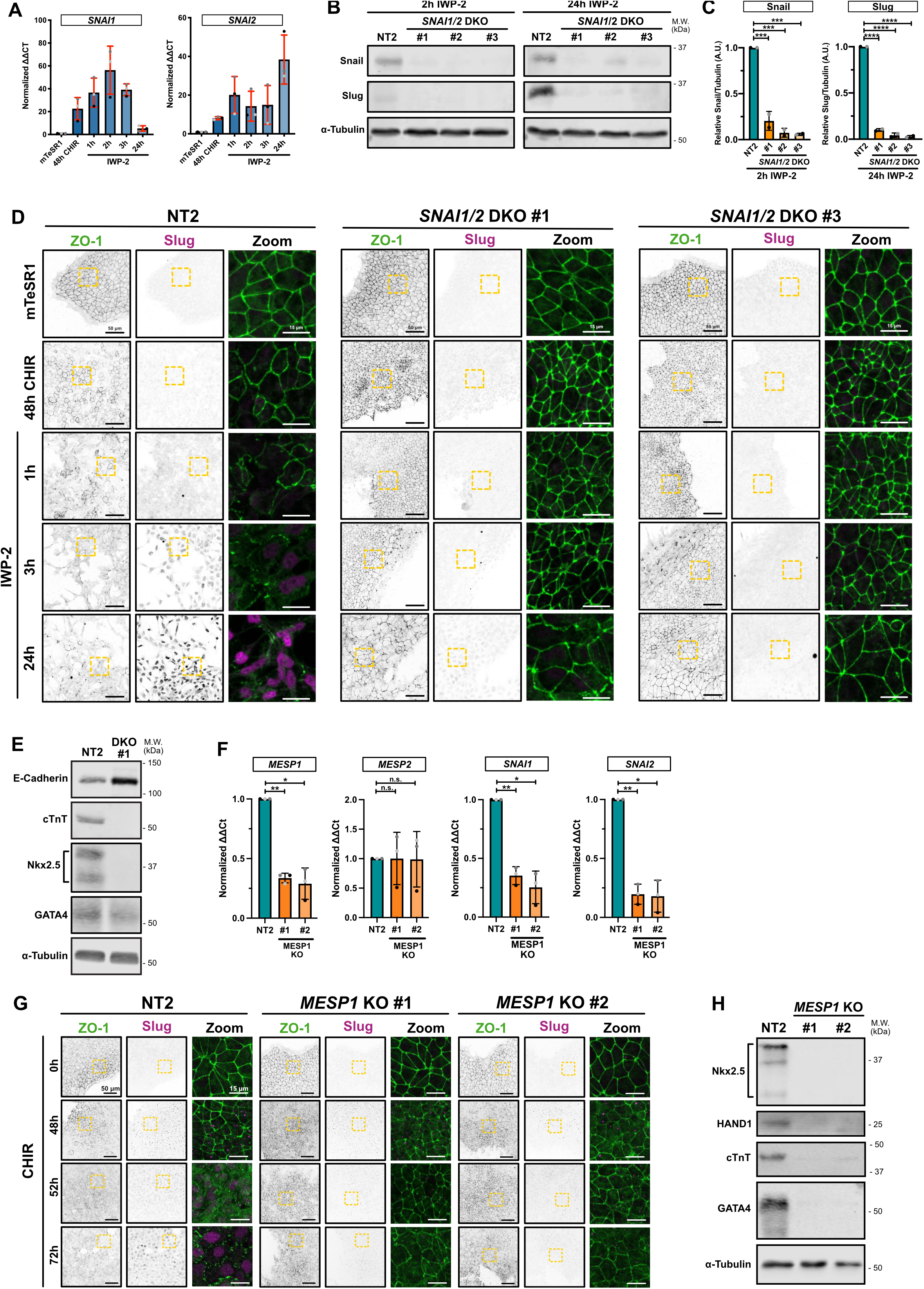
A) qRT-PCR showing expression of *SNAI1* (Snail) and *SNAI2* (Slug) during differentiation. Independent biological repeats are color-coded (n=3). Error bar = Mean +/− S.D. B-C) Immunoblot analysis of three independent *SNAI1*/*SNAI2* double knockout (DKO) hiPSC cell lines (non-clonal). Lysates from Non-targeting (NT2) or DKO cell lines were collected 2h after IWP-2 treatment (to confirm *SNAI1* knockout – Left panel) or 24h after IWP-2 treatment (to confirm *SNAI2* knockout – Right panel). Molecular weights (M.W.) are indicated in kDa (B). Expression levels of 2 independent biological replicates were quantified by densitometry and normalized to NT2. Error bar = Mean +/− S.D. Dunnett’s multiple comparison test was applied (*** p ≤ 0.001, ****p ≤ 0.0001) (C). D) Representative immunofluorescence images of control NT2 and two independent *SNAI1*/*SNAI2* DKO cultures, fixed at different timepoints post-induction of cardiac mesoderm induction and stained for ZO-1 (green) and Slug (Magenta). Maximum intensity projections are shown. Scale bar = 50 μm. Magnified area (yellow dotted square) is shown. Scale bar = 15 μm. E) Immunoblot of control NT2 and *SNAI1*/*SNAI2* DKO #1 hiPSC-derived cardiomyocytes, at 12 days post induction. Expression of EMT marker (E-Cadherin), cardiac lineage markers (cardiac Troponin T, Nkx-2.5, GATA-4) and alpha-Tubulin as loading control were analyzed. Molecular weights (M.W.) are indicated in kDa. F) Relative gene expression of *MESP1*, *MESP2*, *SNAI1* and *SNAI2* obtained from control (NT2) or two independent *MESP1* knockout (#1 and #2) were analyzed by qRT-PCR. ΔΔCt values from RT-PCR were normalized to NT2. Independent biological repeats are color-coded (n=3). Error bar = Mean +/− S.D. Tukey’s multiple comparison test was applied (n.s. p> 0.05, * p ≤ 0.05, ** p ≤ 0.01). G) Representative immunofluorescence images of control NT2 and two independent *MESP1* KO (#1 and #2) non-clonal cell lines, fixed at indicated time, and stained for ZO-1 (green) and Slug (Magenta). Maximum intensity projections are shown. Scale bar = 50 μm. Magnified area (yellow dotted square) is shown as a merge. Scale bar = 15 μm. H) Immunoblot of control NT2 and *MESP1* KO #1 and #2 hiPSC-derived cardiomyocytes, at 12 days post induction. Expression of cardiac lineage markers (cardiac Troponin T, Nkx-2.5, GATA-4, HAND1) and α-Tubulin as loading control were analyzed. Molecular weights (M.W.) are indicated in kDa.

*MESP1* is a pioneer cardiac factor in vivo. During embryonic stem cell differentiation, MESP1 is expressed in ESC-derived cardiac mesoderm progenitors, and is required for cardiac mesoderm specification^14^. Moreover, *MESP1* regulates expression of multiple EMT-promoting genes including *SNAI1*/2. Consistent with these data, we found that CRISPR/Cas9-mediated KO of *MESP1* (Fig 2F) prevented induction of *SNAI1/2* (Fig 2F), blocked the scheduled EMT at 49hrs post-addition of CHIR (Fig 2G), suppressed expression of *NKX2.5, HAND1, cTNT* and *GATA4* (Fig 2H) and prevented differentiation into cardiomyocytes (Supp.Video 3). We conclude that *MESP1* during mesoderm conversion induces *SNAI1/2,* which in turn drive a synchronous EMT that is essential for further differentiation along the cardiac lineage.

### Apoptosis is an essential antecedent to SNAI1/SNAI2 induction and EMT

The initial CHIR treatment of iPSCs caused a rapid, widespread apoptosis, as detected by cleaved caspase 3, PARP cleavage, and Annexin exposure (Fig 1C-H). During this time surviving cells continued to proliferate, increasing the overall cell density (Fig 3A, Supp. Fig S3G-H).

**Figure 3.**
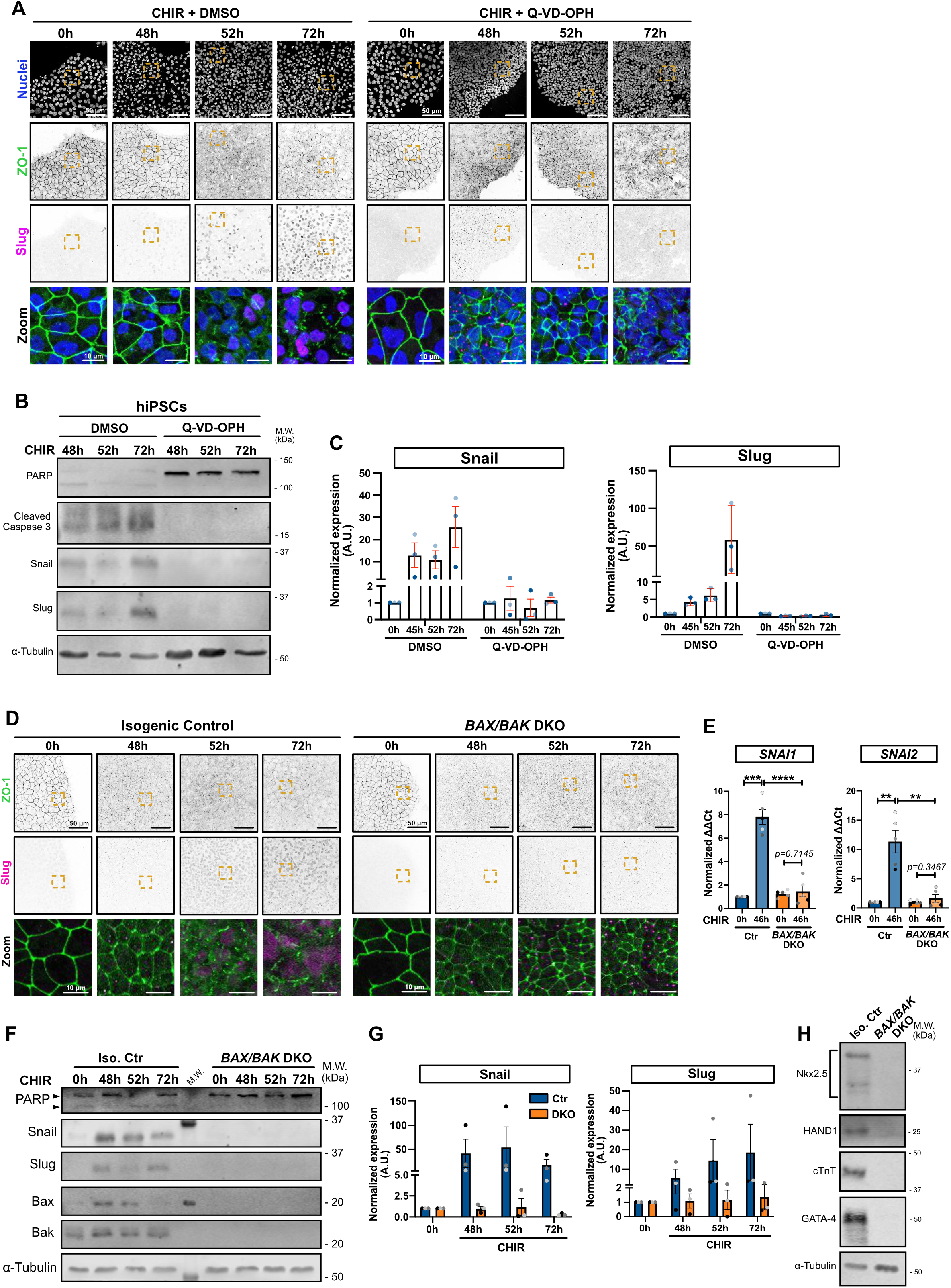
A) Representative immunofluorescence images of hiPSCs co-treated with CHIR and DMSO (left) or CHIR and 10 μM Q-VD-OPH (right) stained for ZO-1 (green) and Slug (magenta) and nuclei (blue). Maximum intensity projections are shown. Scale bar = 50 μm. Magnified area (yellow dotted square) is shown as a merge. Scale bar = 10 μm. B-C) Immunoblot analysis of hiPSCs co-treated with CHIR plus DMSO or 10 μM Q-VD-OPH. Molecular weights (M.W.) are indicated in kDa (B). Normalized expression of Snail and Slug was quantified by densitometry across 3 independent biological replicates (color-coded). Mean +/− S.E.M. (C). D) Representative immunofluorescence images of isogenic control and *BAX/BAK* double knock-out (DKO) hiPSCs, treated with CHIR and stained for ZO-1 (green) and Slug (magenta). Maximum intensity projections are shown. Scale bar = 50 μm. Magnified area (yellow dotted square) is shown as a merge. Scale bar = 15 μm. E) Relative gene expression of *SNAI1* and *SNAI2* obtained from isogenic control (Ctr) or *BAX/BAK* DKO hiPSCs, induced or not with CHIR for 46h and analyzed by qRT-PCR. ΔΔCt values were normalized to un-induced control cells. Independent biological repeats are color-coded (n=5). Error bar = Mean +/− S.E.M. Paired t-test was applied to compare 0h vs 46h and unpaired t-test was applied to compare Ctr vs DKO (** p ≤ 0.01, *** p ≤ 0.001, **** p ≤ 0.0001). F-G) Immunoblot analysis of isogenic control (Iso. Ctr) and *BAX/BAK* DKO hiPSCs, treated with CHIR. Molecular weights (M.W.) are indicated in kDa (F). Normalized expression of Snail and Slug was quantified by densitometry across 3 independent biological replicates (color-coded). Mean +/− S.E.M. (G). H) Immunoblot of isogenic control (Iso. Ctr) and *BAX/BAK* DKO hiPSC-derived cardiomyocytes, at 12 d post induction. Expression of cardiac lineage markers (cardiac Troponin T, Nkx-2.5, GATA-4, HAND1) and α-Tubulin as loading control were analyzed.

To test whether apoptosis has any impact on EMT and conversion along the cardiac mesoderm lineage, we blocked cell death in cultures treated with CHIR, using the pan-caspase inhibitor Q-VD-OPH. Remarkably, this drug totally prevented the scheduled EMT (Fig. 3A). The cells maintained a cobblestone appearance, retained tight junctions, grew to high density (Supp. Fig S3G), and failed to express Snail or Slug (Fig 3A-C). Inhibition of apoptosis was marked by loss of cleaved Caspase 3 and PARP (Fig 3B). A similar effect of Q-VD-OPH on EMT was observed in human ESCs (Supp. Fig S3A-C).

To validate the connection to cell death we generated iPSC lines deleted for the apoptotic executioner proteases Caspase 3 or Caspase 9, using CRISPR/Cas9-mediated gene editing (Supp. Fig S3D). These lines did not proliferate as rapidly as the parental WT or the control non-target (NT) iPSCs, but nonetheless formed island cultures. Notably, treatment with CHIR did not cause TJ disassembly, even after prolonged incubation, and did not induce *SNAI1/2* (Supp. Fig 3E-F).

Next, to prevent the apoptotic events upstream of caspase activation, we deleted the pro-apoptotic genes *BAX* and *BAK*^15^. Unlike the Caspase 3 and 9 KO lines, these double knockout (DKO) cells grew at rates comparable to control cells (Supp. Fig S3H), but again, no EMT was detected after treatment with CHIR as determined by persistence of TJs (Fig 3D) and by the absence of Snail and Slug expression at both the protein and mRNA levels (Fig 3E – G).

Moreover, cardiomyocyte differentiation was also blocked (Fig 3H; Supp. Fig 3I and Supp.Video 4). Similarly, addition of Q-VD-OPH during both, CHIR-99021 and IWP-2 steps, blocks induction of the cardiac marker HAND1 (Supp. Fig S3J).

Together, these data strongly support an unanticipated requirement for apoptosis, induced after WNT activation, to permit the entry of pluripotent cells into the mesoderm lineage. We note that cell death has been observed in early mouse embryos, just prior to gastrulation and in primitive streak, although its function is unknown^16^.

### Apoptosis is required very early in selection of the mesoderm lineage

To determine at which stage apoptosis permits pluripotent stem cells to differentiate towards cardiomyocytes, we treated *BAX/BAK* DKO iPSCs with CHIR, then harvested cells at 46 hrs for QRT-PCR. Surprisingly, even very early changes in gene expression, including *TBXT* (*T/Bra)* and *EOMES* were drastically reduced or suppressed. A similar suppression was caused by the treatment of WT iPSCs with QVD-OPH prior to addition of CHIR (Fig 4B). The failure to express these genes was confirmed by immunofluorescence (Fig. 4C) and by immunoblotting for T/Bra (Fig. 4D). Later changes associated with mesoderm induction, including *TBX6* and *MESP1* were also inhibited (Fig. 4A-B); however, expression of the pluripotent marker *Nanog* decreased on schedule (Fig 4E). Importantly, DKO iPSCs lacking BAX and BAK can still enter the neuronal lineage^15^.

**Figure 4.**
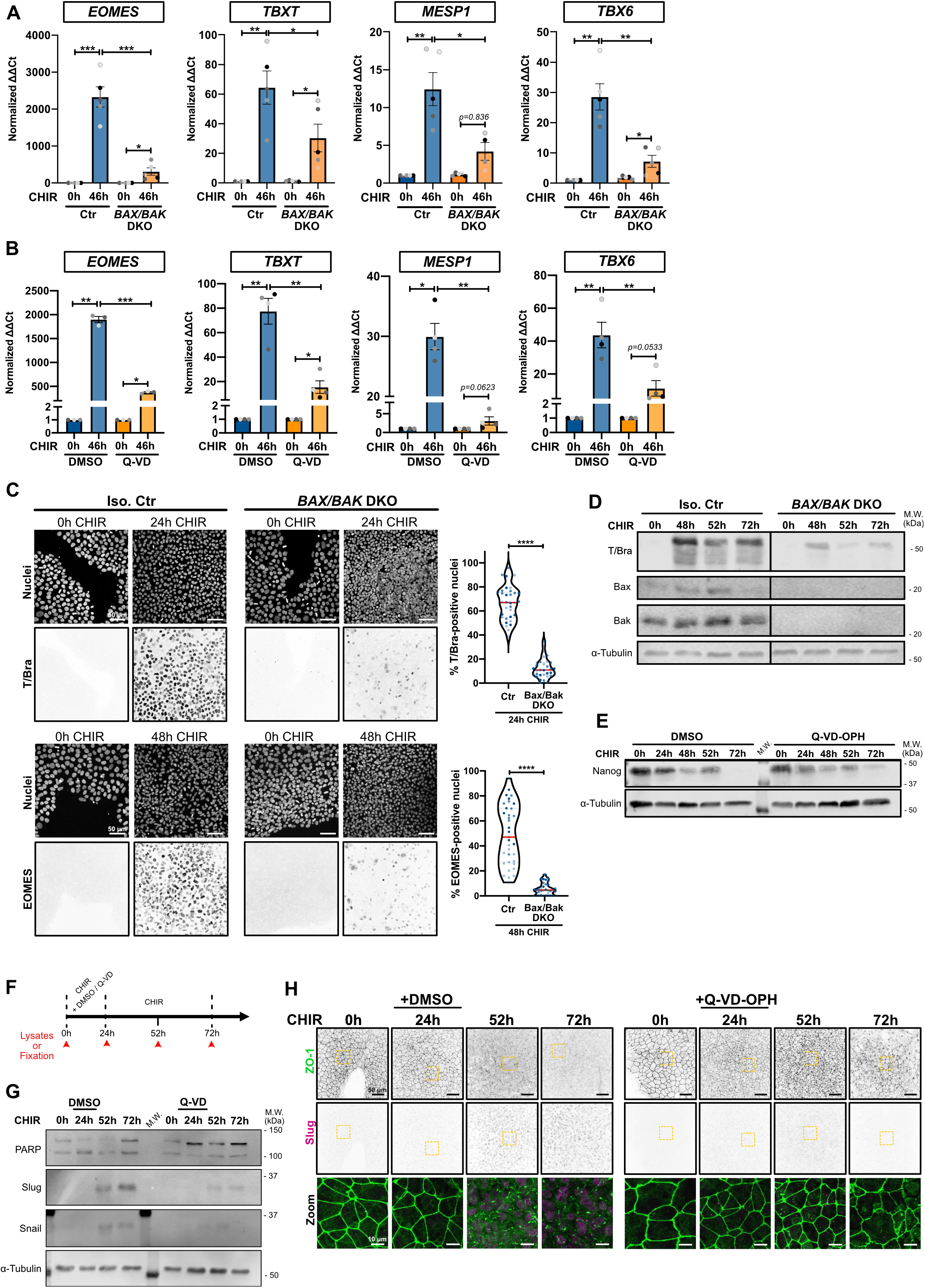
A)Relative gene expression of *EOMES, TBXT, MESP1, TBX6* obtained from isogenic control (Ctr) or *BAX/BAK* DKO hiPSCs, induced or not with CHIR for 46h and analyzed by qRT-PCR. ΔΔCt values were normalized to un-induced cells. Independent biological repeats are color-coded (n=5). Error bar = Mean +/− S.E.M. Paired t-test was applied to compare 0h vs 46h and unpaired t-test was applied to compare Ctr vs DKO (* p ≤ 0.05, ** p ≤ 0.01, *** p ≤ 0.001). B) Relative gene expression of *EOMES, TBXT, MESP1, TBX6* obtained from WT hiPSCs, induced or not with CHIR and DMSO or CHIR and Q-VD-OPH for 48h and analyzed by qRT-PCR. ΔΔCt values were normalized to un-induced cells. Independent biological repeats are color-coded (n=3-4). Error bar = Mean +/− S.E.M. Paired t-test was applied to compare 0h vs 48h and unpaired t-test was applied to compare DMSO vs Q-VD (* p ≤ 0.05, ** p ≤ 0.01, *** p ≤ 0.001). C) Representative immunofluorescence images of isogenic control (Iso. Ctr) and *BAX/BAK* DKO hiPSCs stained for T/Bra (top) and EOMES (bottom) and nuclei, before (0h) and after CHIR treatment. Maximum intensity projections are shown. Scale bar = 50 μm. Violin plots summarize quantification of the percentage of T/Bra-positive (top) and EOMES-positive (bottom) nuclei after CHIR treatment (Median: plain red line – Quartiles: black dotted lines). Independent biological repeats are color-coded (n=3, 10-12 random fields of view/repeat). Mann-Whitney test was applied (**** p ≤ 0.001). D) Immunoblot of isogenic control (Iso. Ctr) and *BAX/BAK* DKO hiPSCs analyzing T/Bra expression following CHIR addition. Molecular weights (M.W.) are indicated in kDa. E) Immunoblot of WT iPSC treated with CHIR +/− 10 μM Q-VD-OPH and analyzed for Nanog expression. Molecular weights (M.W.) are indicated in kDa. F) Treatment schedule is indicated and timepoints for lysate collection and fixation is depicted by red arrows. G-H) Immunoblot analysis (G) and representative immunofluorescence images (H) of hiPSCs treated with CHIR +/− 10 μM Q-VD-OPH for 24h, then Q-VD-OPH was washed out and cells were incubated for another 28h or 48h with CHIR (respectively 52h and 72h timepoint).

These data suggest that, even though apoptosis continues to occur for >40 hours after activation of WNT signaling, it is necessary for a very early event following initiation of mesoderm specification. To further test this hypothesis, we treated WT iPSCs with CHIR and added Q-VD-OPH only for the first 24 hrs, after which time the medium was replaced to CHIR alone for an additional 48 hrs (Fig 4F) or, as a control, iPSCs were treated with CHIR alone until 24 hrs had passed, then incubated with CHIR + Q-VD-OPH (Supp. Fig S4C). Notably, early addition of this inhibitor completely blocked the subsequent EMT and strongly reduced Snail/Slug induction (Fig. 4G-H); but addition after 24 hrs had no effect and EMT occurred on schedule (Supp. Fig S4C-D)). Together, these data highlight an unanticipated essential role for apoptosis in the initial steps towards mesoderm specification.

### A soluble factor from apoptotic cells permits pluripotent cells to enter the mesoderm lineage

How might apoptosis allow pluripotent stem cells to enter the cardiac mesoderm lineage? We considered two possibilities. First, cell death might relax the space constraints between cells in a colony, allowing them to stretch and generate tension, which in principle could activate signaling through YAP/TAZ or some other pathway to promote mesoderm conversion.

Alternatively, apoptotic cells might release a soluble factor that promotes mesoderm conversion. We discounted the first hypothesis as unlikely, because we did not detect any increase in nuclear YAP localization after CHIR treatment (Supp. Fig S5A). To test the second hypothesis, we treated both WT and the *BAK/BAX* DKO iPSCs for 24 hrs with CHIR then replaced the DKO medium with conditioned medium (CM) from the apoptosing WT iPSCs, and continued to incubate the DKO cells for a further 48 hrs (Fig 5A). Remarkably, the DKO cells receiving the CM underwent a dramatic EMT (Fig 5B). To test if the CM also relieved the blockade to expression of mesoderm lineage genes caused by the inability of the DKO cells to undergo apoptosis, we performed RT-qPCR and found significant increases in *EOMES, MESP1, TBX6,* and *SNAI1/2* (Fig 5C). These results clearly demonstrate that a soluble factor released by WNT-induced apoptosis of iPSCs is required for mesoderm conversion and consequent EMT of the surviving iPSC population.

**Figure 5.**
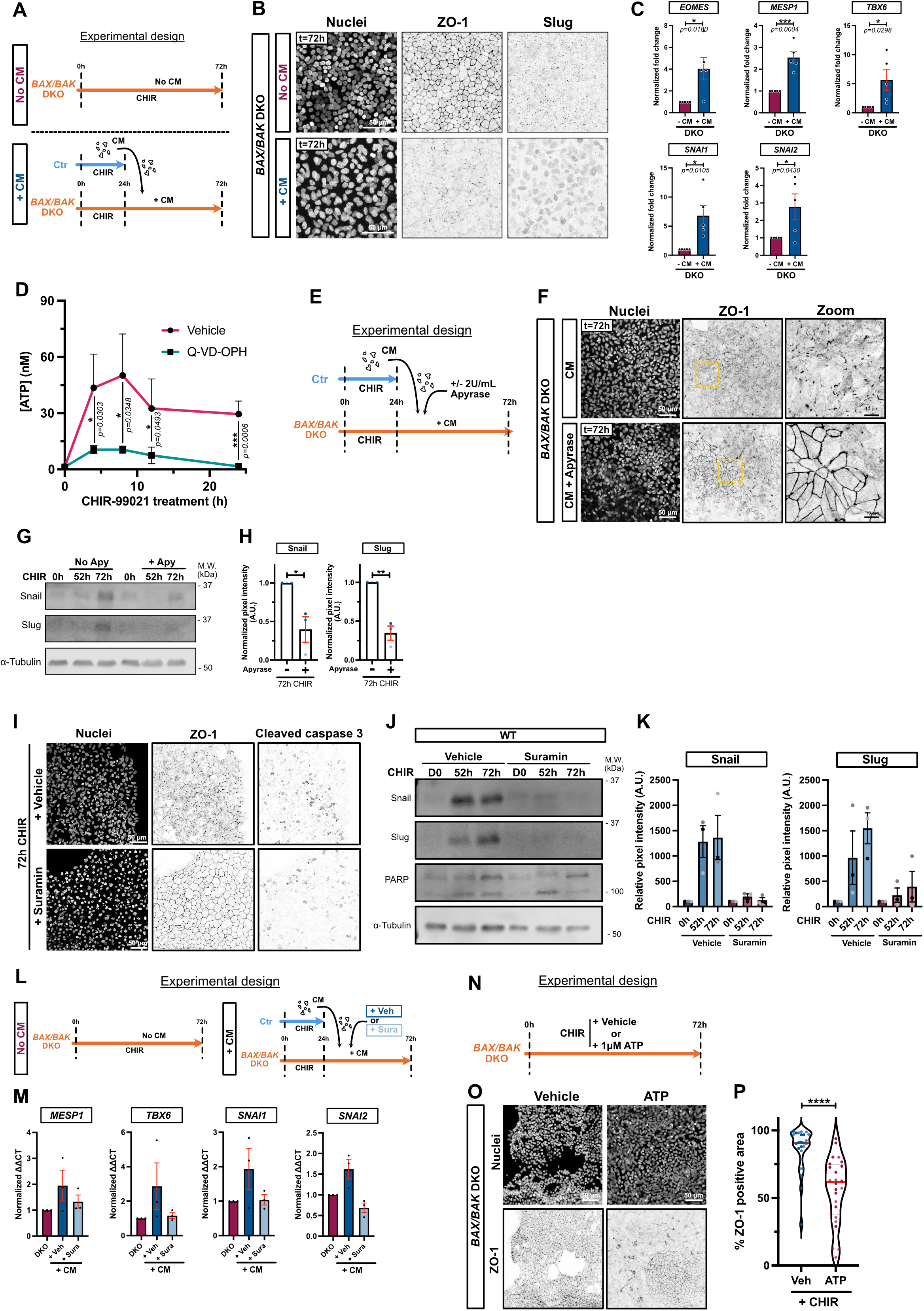
A)Timeline of conditioned media (CM) experiment. B) Immunofluorescence pictures of *BAX/BAK* DKO cells, pre-treated with CHIR for 24h and incubated for another 48h without (top) or with (bottom) condition media. Cells were fixed at 72h and stained for ZO-1, Slug, and nuclei. Maximum intensity projections are shown. Scale bar = 50 μm. C) Relative gene expression of *EOMES, MESP1, TBX6, SNAI1 and SNAI2* obtained from *BAX/BAK* DKO hiPSCs treated as described in A and analyzed by qRT-PCR across 5 independent biological repeats. Fold change in ΔΔCt values was normalized to - CM condition. Error bar = Mean +/− S.E.M. Paired t test was applied (* p ≤ 0.05, *** p ≤ 0.001). D) Time course of ATP release from dying cells. Luciferase assay was performed on supernatants from hiPSCs treated with CHIR +/− 10 μM Q-VD-OPH at indicated timepoints. Luminescence was converted to [ATP] using a standard curve. Three independent biological replicates (2 technical replicates each) were plotted as a line graph with mean +/− S.D. Two-way ANOVA with Sidak’s multiple comparisons test was applied (* p ≤ 0.05, *** p ≤ 0.001). E) Timeline of apyrase-treated conditioned media (CM) experiment. F) Representative immunofluorescence pictures of *BAX/BAK* DKO cells fixed after adding apyrase-treated CM as depicted in (E). Cells were stained for ZO-1 and nuclei. Maximum intensity projections are shown. Scale bar = 50 μm. Magnified area (yellow dotted square) is shown for the ZO-1 channel. Scale bar = 10 μm. G-H) Immunoblot analysis of WT hiPSCs treated with CHIR +/− 2U/mL Apyrase and probed for Snail, Slug and alpha-Tubulin (G). Normalized expression of Snail and Slug was quantified by densitometry across 3 independent biological replicates (color-coded). Mean +/− S.E.M. Unpaired t-test was applied (* p ≤ 0.05, ** p ≤ 0.01) (H). I) Immunofluorescence pictures of WT hiPSCs treated with CHIR +/− 100 μM Suramin and stained for ZO-1, cleaved-caspase 3 and DNAi. Maximum intensity projections are shown. Scale bar = 50 μm. J-K) Immunoblot analysis of WT hiPSCs treated with CHIR +/− 100 μM Suramin and probed for Snail, Slug, PARP and α-Tubulin (J). Normalized expression of Snail and Slug was quantified by densitometry across 3 independent biological replicates (color-coded) (K). L) Timeline of conditioned media (CM) experiment with Suramin. M) Relative gene expression of *MESP1, TBX6, SNAI1 and SNAI2* obtained from *BAX/BAK* DKO hiPSCs treated as described in K and analyzed by qRT-PCR across 3 independent biological repeats. Mean +/− S.E.M. N) Timeline of ATP sufficiency experiment O) Immunofluorescence images of *BAX/BAK DKO* hiPSCs treated with CHIR +/− 1 μM ATP and stained for ZO-1 and nuclei. Maximum intensity projections are shown. Scale bar = 50 μm. P) Violin plot showing percentage of ZO-1 positive areas across 2 independent biological samples (3 large images per repeat (5mm × 5mm) obtained by tlling). Mann-Whitney test was applied (**** p ≤ 0.001).

### ATP provides an essential signal through the purinergic P2Y receptor

Apoptotic cells generate soluble find-me signals that recruit macrophages and membrane-associated eat-me signals to promote engulfment^17,18^. Nucleotides (ATP, UTP) have been identified as potent find-me signals^19^. We first tested, using a luciferase assay, whether apoptosing iPSCs release ATP. Treatment with CHIR caused a significant increase in extracellular ATP within 8 hrs (Fig 5D). Moreover, treatment with apyrase, to hydrolyze nucleotides, partially blunted the EMT induced by CM on *BAX/BAK* DKO (Fig. 5E-F). This result was confirmed in a setting where apoptosis was blocked in WT cells using Q-VD-OPH for 24hrs prior to adding the CM +/− apyrase (Supp. Fig S5B-C). Moreover, WT hiPSCs co-treated with CHIR and apyrase showed a two-fold reduction in Snail and Slug expression (Fig 5G, H).

Purinergic P2Y receptors bind ATP among other nucleotides, and function as chemo-attractants for macrophages^19,20^. Strikingly, the P2 receptor inhibitor suramin totally blocked EMT in WT iPSCs treated with CHIR, even though many of these cells still underwent apoptosis (Fig 5I). Snail and Slug were also completely suppressed (Fig. 5J-K). Additionally, suramin blocked the effect of CM on *BAK/BAX* DKO cells, preventing EMT and accompanying gene expression changes (Fig 5L-M).

Finally, addition of ATP to *BAK/BAX* DKO cells treated with CHIR also induced a partial EMT (Fig 5N-P). However, induction of apoptosis by UV irradiation, in the absence of WNT signaling, did not cause EMT (Supp. Fig. S5D-E).

We conclude, therefore, that WNT signaling induces two distinct and complementary responses in pluripotent stem cells, both of which are needed for commitment to cardiogenesis: the first activating early apoptosis, triggering the release of nucleotides, including ATP, which through P2Y receptor engagement act in a paracrine fashion to permit the stem cells to enter the mesoderm lineage; while the second drives differentiation of the responsive cells through primitive streak towards cardiac mesoderm (Supp. Fig S5F).

## Discussion

Apoptotic cells release find-me and eat-me signals that ensure their rapid clearance from tissues by macrophages and other phagocytic cells^7,18^. Find-me signals include several molecules, including nucleotides that are recognized by purinergic G-protein coupled P2Y receptors and act as chemo-attractants^9^. P2Y agonists have multiple biological functions in addition to apoptotic clearance^20^ but have not previously been implicated in early developmental decisions. Our discovery that suppression of apoptosis in human iPSCs and ESCs completely blocks specification along the mesoderm lineage in response to WNT activation was, therefore, highly unexpected. Even early changes in gene expression, such as the induction of EOMES, are prevented, resulting in a later block in *SNAI1/2* expression and in the subsequent EMT, which we showed is essential for cardiac mesoderm commitment. This blockade must, however, occur after escape from pluripotency, because the drop in Nanog expression induced by WNT activation occurred normally. Moreover, *BAK/BAX* double KO iPSCs can still successfully enter the neural tube lineage^15^. A previous report identified caspase activity as being required for the differentiation of ESCs in response to retinoic acid^13^. However, the mechanism in this case appears to be through caspase-induced cleavage and deactivation of Nanog, rather than through the generation of a soluble paracrine signal. Apoptosis occurs pre-gastrulation during mouse embryogenesis^16^, but apoptosis-defective mice generally progress through embryogenesis. *CASP3* /*CASP7* double KO mice die perinatally from cardiovascular defects^21^. A triple KO mouse line lacking BAX, BAK and BOK also develops through embryogenesis^22^. This observation emphasizes the idea that other types of cell death (such as ferroptosis) might also occur in embryogenesis to release nucleotides; additionally, nucleotides can be released through pannexin 1 channels in response to multiple other stresses, which might provide the necessary signal in embryogenesis^20,23^. Finally, we noted that ATP was insufficient to trigger 100% of the cells to undergo EMT; moreover, the conversion occurs in clusters across the cell colonies. It is possible that additional metabolites released by apoptosing cells contribute to the signal, or that the nucleotides are degraded before they can trigger differentiation of the entire cell population. The patchiness of the response suggests cell-cell communication might promote the EMT, or that groups of cells are in different initial states that are less or more susceptible to purinergic signaling.

Overall, it is remarkable that the death of a fraction of pluripotent stem cells is required for differentiation of the survivors, through a paracrine find-me signal that usually functions for apoptotic cell clearance. It will be of interest to determine if other developmental processes require similar signaling mechanisms.

## Material & Methods

### Reagents

Common lab reagents are listed in **Table 1**.

**Table 1:**
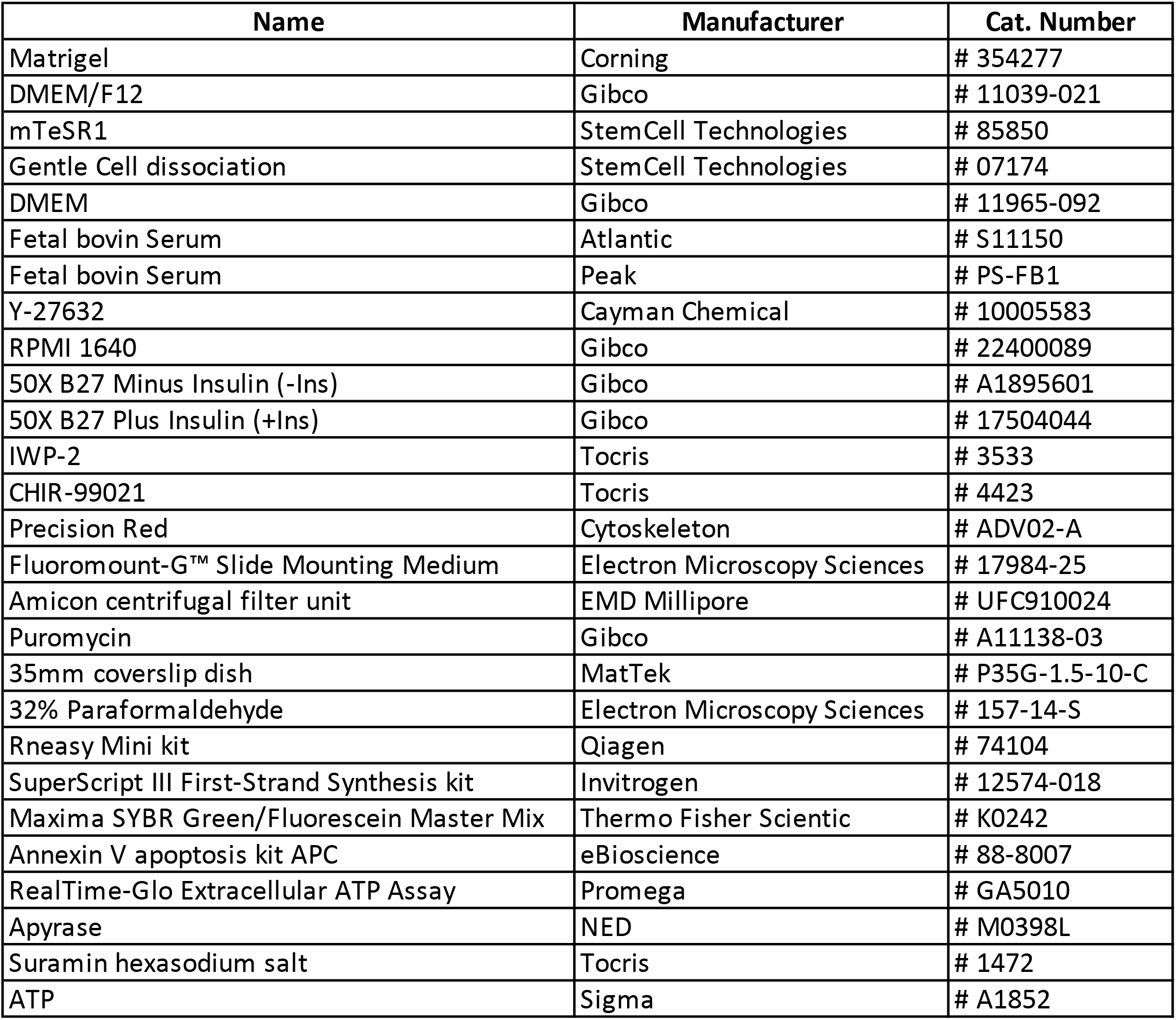
Lab reagents

### Cell lines and Cell culture

GM25256 hiPSCs were obtained from the Coriell Institute and were derived from a healthy 30-year-old male. mEGFP-TJP1 knock-in GM25256 hiPSC cell line was obtained from the Allen Cell Collection, Coriell Institute (Cell ID AICS-0023 cl.20). *BAX/BAK* double knock-out hiPSC GM25256 cell line were obtained from Dr. Vivian Gama (Joshi et al, https://doi.org/10.1038/s41419-020-03002-x). Human embryonic stem cell line H9 (WA09) was obtained from WiCell Research Institute (Wisconsin). All experiments using hESCs were performed using the WA09 (H9) cell line under the supervision of the Vanderbilt Institutional Human Pluripotent Cell Research Oversight (VIHPCRO) Committee (Protocol IRB # 160146 to VG).

hiPSCs and hESCs H9 cell lines were cultured on Matrigel coated 6-well plates (Matrigel diluted at 42 μg/mL in DMEM/F12 media) and grown in mTeSR1 medium. Medium was changed daily until cells reached 70% confluence. Cells were passaged using Gentle cell dissociation reagent for 4 min, resuspended in mTseR1 as small clusters and replated at 1:7.

HEK-293T cells were obtained from ATCC and maintained in Dulbecco’s modified Eagle’s medium supplemented with 10% fetal bovine serum and passaged at 1:10 every 2-3 d. All cell lines used in this study were maintained at 37 °C and under 5% CO_2_.

### Cell freezing and thawing

hiPSCs and hESCs were collected as previously described and centrifuged at 1000 rpm for 3 min. Pellets were resuspended in mTeSR1 supplemented with 10% DMSO and aliquoted in cryovials. Cells were first transferred at −80 °C for 24h before long-term storage in liquid nitrogen. hiPSCs and hESCs were slowly thawed using mTeSR1 media, centrifuged and resuspended in mTeSR1 supplemented with 10 μM of Y-27632 for 24h.

### Cardiomyocyte differentiation protocol

This protocol was adapted from Lian et al^2,24^ (GiWi protocol).

Briefly once confluency reached 70-80%, cells were treated with RPMI 1640 supplemented with 1X B27 minus Insulin and 7.5 μM CHIR-99021 for 48h. At 48h after CHIR addition, media was aspirated and replaced with RPMI 1640, supplemented with 1X B27 minus Insulin and 7.5 μM of IWP-2 for 48h. Then, cells were incubated for 48h with RPMI 1640/1X B27 minus Insulin, before maintaining them in RPMI 1640/1X B27 plus insulin every 3 d. Spontaneous and homogenous beating should be observed within 10-12 days after the protocol initiation.

HiPSC-derived cardiomyocytes (hiPSC-CMs) used in Figure 5G were generated using the small molecules CHIR-99021 (Selleck Chemicals) and IWR-1 (Sigma). Cardiac differentiation media were defined as M1 (RPMI 1640 with glucose with B27 minus insulin), M2 (RPMI 1640 minus glucose with B27 minus insulin), and M3 (RPMI 1640 with glucose with B27). When hiPSCs reached 60% confluence, cardiac differentiation was initiated (day 0). At d 0, hiPSCs were supplemented in M1 with 6 μM CHIR-99021. On d 2, the media was changed to M1. On d 3, cells were treated with 5 μM IWR-1 in M1. Metabolic selection was started at day 10 and cells were treated with M2 from d 10 to 16. On d 16, cells were transitioned to M3. Media was changed every other day until d 30.

### SDS-PAGE and Western blotting

Cells were washed in 1X PBS. Lysates were obtained by scraping cells in lysis buffer (150 mM NaCl, 10 mM Tris-HCl pH 7.5, 1 mM ethylenediaminetetraacetic acid (EDTA), 1% Triton X-100, 0.1% SDS, 1X protease and phosphatase inhibitors) followed by a 5 min incubation on ice and centrifugation at 16000 rpm for 10 min at 4°C. Protein concentration was measuring using Precision Red.

30 μg of proteins were resolved on bis-tris acrylamide gels and transferred onto nitrocellulose membrane for 90min at 110V. Membranes were blocked for 30 min in 5% non-fat milk in TBS-T (10 mM Tris pH 8.0, 150 mM NaCl, 0.5% Tween 20) before overnight incubation with primary antibodies (**Table 2**) at 4°C with gentle rocking. Membranes were washed in TBS-T and incubated 1h at room temperature with Alexa-Fluor conjugated secondary antibodies (**Table 2**). Membranes were washed in TBS-T and scanned using the LI-COR Odyssey CLx.

**Table 2:** Antibodies

| Name | Manufacturer | Cat. Number | Application | Dilution |
| --- | --- | --- | --- | --- |
| E_Cadherin | Cell Signaling Technologies | # 3195 | WB | 1:1000 |
|  | Invitrogen | # 13-1900 | IF | 1:500 |
| Slug | Cell Signaling Technologies | # 9585 | WB | 1:1000 |
|  |  |  | IF | 1:500 |
| Snail | Cell Signaling Technologies | # 3895 | WB | 1:1000 |
| Vimentin | Cell Signaling Technologies | # 5741 | IF | 1:500 |
| Zeb1 | Cell Signaling Technologies | # 3396 | WB | 1:1000 |
| GATA4 | Cell Signaling Technologies | # 36966 | WB | 1:1000 |
| Nanog | Cell Signaling Technologies | # 4903 | WB | 1:1000 |
| Oct4 | Cell Signaling Technologies | # 75463 | WB | 1:1000 |
| Sox2 | Cell Signaling Technologies | # 3579 | WB | 1:1000 |
| Alpha-Tubulin | Sigma-Aldrich | # T9026 | WB | 1:3000 |
| ZO-1 | Invitrogen | # 339100 | WB | 1:200 |
|  |  |  | WB | 1:500 |
| cTnT | Gift from Prof. Dylan Burnette |  | WB | 1:500 |
| Nkx2.5 | Cell Signaling Technologies | # 8792 | WB | 1:1000 |
| Cleaved caspase 3 | Cell Signaling Technologies | # 9664 | WB | 1:1000 |
|  |  |  | IF | 1:200 |
| PARP | Cell Signaling Technologies | # 9542 | WB | 1:1000 |
| Scribble | Santa Cruz Biotechnology | # sc-11048 | WB | 1:500 |
|  |  |  | IF | 1:200 |
| LLGL2 | Santa Cruz Biotechnology | # sc-130158 | WB | 1:500 |
| PAR-3 | Homemade |  | WB | 1:500 |
|  |  |  | IF | 1:200 |
| PKC-ζ | Santa Cruz Biotechnology | # sc-17781 | WB | 1:500 |
|  |  |  | IF | 1:200 |
| PALS-1 | Homemade |  | WB | 1:500 |
|  |  |  | IF | 1:200 |
| HAND-1 | R & D Systems | # AF3168 | WB | 1:500 |
| EOMES | Cell Signaling Technologies | # 81493 | IF | 1:200 |
| T/Bra | abcam | # ab209665 | IF | 1:200 |
|  |  |  | WB | 1:1000 |
| alpha actinin 2 | Signa | A7811 | IF | 1:500 |
| YAP | Novus | # NB110-58358 | IF | 1:200 |

**Table 2:** Secondary Antibodies

|  |  |  |  |  |
| --- | --- | --- | --- | --- |
| Alexa Fluor 680 donkey anti-mouse IgG (H+L) | Invitrogen | # A10038 | WB | 1:10000 |
| Goat anti-Rabbit (H&L), DyLight 800 4X PEG conjugate | Invitrogen | # SA535571 | WB | 1:10000 |
| Alexa Fluor 594 goat anti-rabbit IgG (H+L) | Invitrogen | # A11037 | IF | 1:500 |
| Alexa Fluor 488 goat anti-mouse IgG (H+L) | Invitrogen | # A11029 | IF | 1:500 |
| Alexa Fluor 488 goat anti-rat IgG (H+L) | Invitrogen | # A11006 | IF | 1:500 |

**Table 2:** Others

|  |  |  |  |  |
| --- | --- | --- | --- | --- |
| Alexa Fluor 647 Phalloidin | Invitrogen | # A22287 | IF | 1:500 |
| Hoechst 33342 | Invitrogen | # H1399 | IF | 1:500 |

All images were analyzed using Image Studio Lite v. 5.2.5.

### Immunofluorescence

Cells were grown on Matrigel-coated coverslip, fixed with 4% paraformaldehyde for 10 min, permeabilized (20 mM glycine, 0.05% Triton X-100) for 10 min and blocked with 5% BSA-PBS for 30 min. Primary and secondary antibodies were diluted in blocking buffer and incubated for 1 h in a dark, humidified chamber. Coverslips were washed three times in PBS before being mounted on glass slides using Fluoromount-G™ Slide Mounting Medium.

Images were taken using an inverted Nikon A1-R confocal microscope equipped with a 40x oil objective (NA 1.2). 0.5 μm Z-stack covering the entire cell height were obtained.

Super-resolution images for Figure 5G were acquired using a Nikon SIM microscope equipped with a 1.49 NA 100x Oil objective an Andor DU-897 EMCCD camera.

Images were processed and analyzed using Fiji software (ImageJ version 2.1.0/1.53c).

### Live Cell Imaging

mEGFP-TJP1 hiPSCs were plated on Matrigel-coated MaTek 35mm dishes. Cells were imaged every 10-15 min on a Nikon A1-R with a 40X oil objective (NA 1.2) and equipped with a heated CO2 chamber. 2-3 μm Z-stack were obtained and images were processed and analyzed using Fiji software (ImageJ version 2.1.0/1.53c).

### Generation of knock-out cell lines

Single-guide RNA was selected using ChopChop^25^ and Benchling design tools and are listed in **Table 3**. Annealed oligonucleotides were cloned into pLentiCrispRv2-Puro as described by Sanjana et al (10.1038/nmeth.3047). HEK-293T cells were seeded on 15 cm dish to 50% confluence and transfected using calcium phosphate. Briefly, 50 μg of the lentiviral plasmid, 37.5 μg of pSPAX2 (Addgene 8454) and 15 μg of pMD2G (Addgene 12260) were combined to 1125 μl of sterile water, complemented with 125 μl of 2.5M CaCl_2_. While vortexing, 1.25 ml of filter sterilized 2× HEPES-buffered saline (50 mM Hepes, 10 mM KCl, 12 mM Dextrose, 280 mM NaCl, 1.5 mM Na_2_PO_4_, pH 7.04) was added, and the solution was incubated 5 min at RT before adding to HEK-293T cells. Medium was removed after 6-8 h and replaced with 15 ml of 10% FBS DMEM. Lentiviruses were collected after 48h, concentrated using Amicon centrifugal filter units (100 kDa cut-off) and stored at −80°C. hiPSCs were transduced in suspension for 24h and then selected using 1 μg/mL Puromycin.

**Table 3:**
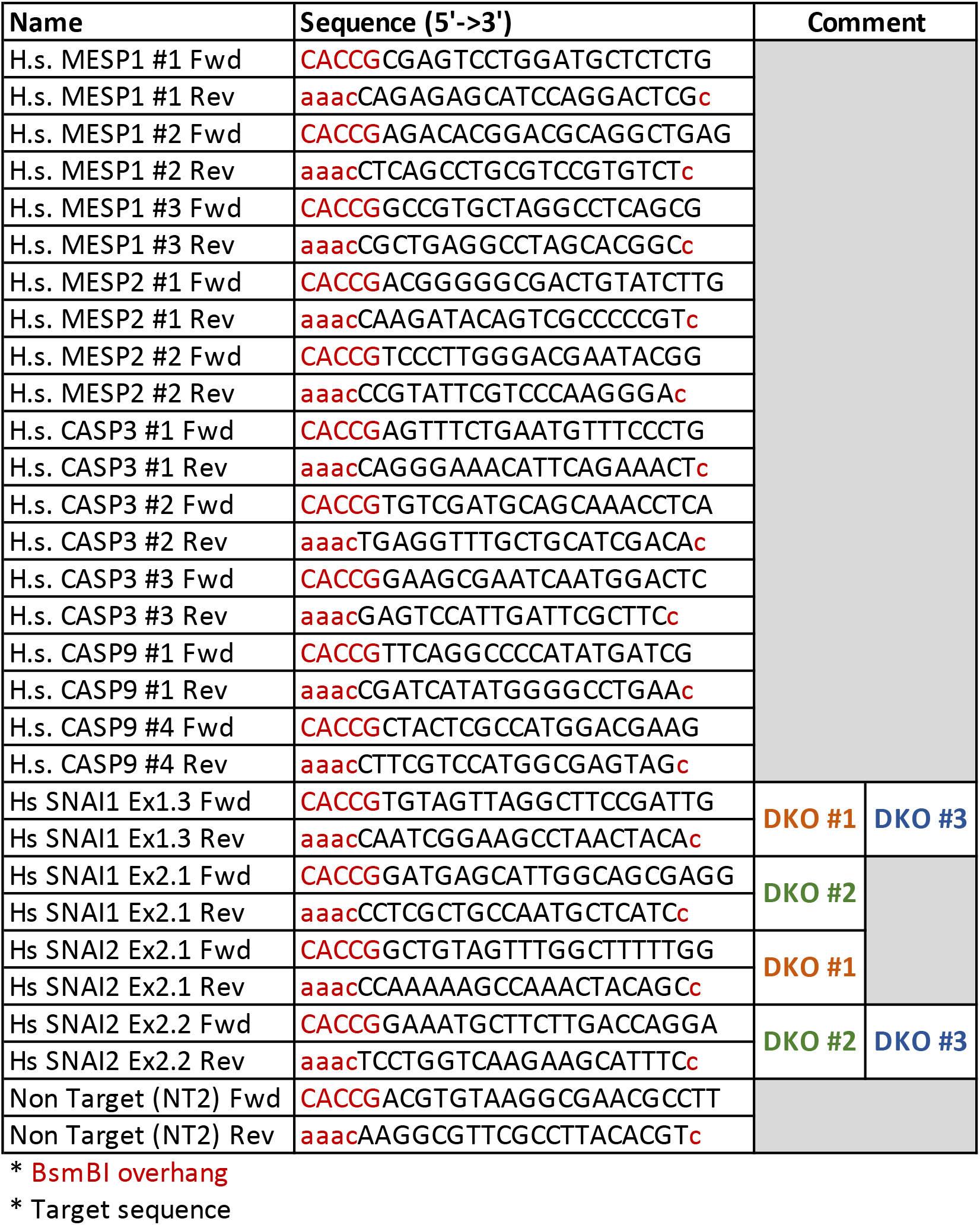
CRISPR-Cas9 sgRNA *

### RNA isolation and RT-qPCR

RNA was isolated using RNeasy Mini kit. 1 μg of RNA was reverse transcribed to cDNA using SuperScript III First-Strand Synthesis System and diluted 1:10 in water. 4.5 μL of cDNA was mixed with 7.5 μL Maxima SYBR Green/Fluorescein Master Mix and 3 μL of primers (1 mM each) (**Table 4**). qPCR was performed on a BioRad CFX96 Thermocycler and Ct values from technical triplicates were average and used to calculate the relative gene expression normalized to GAPDH, using the ΔΔCt formula.

**Table 4:**
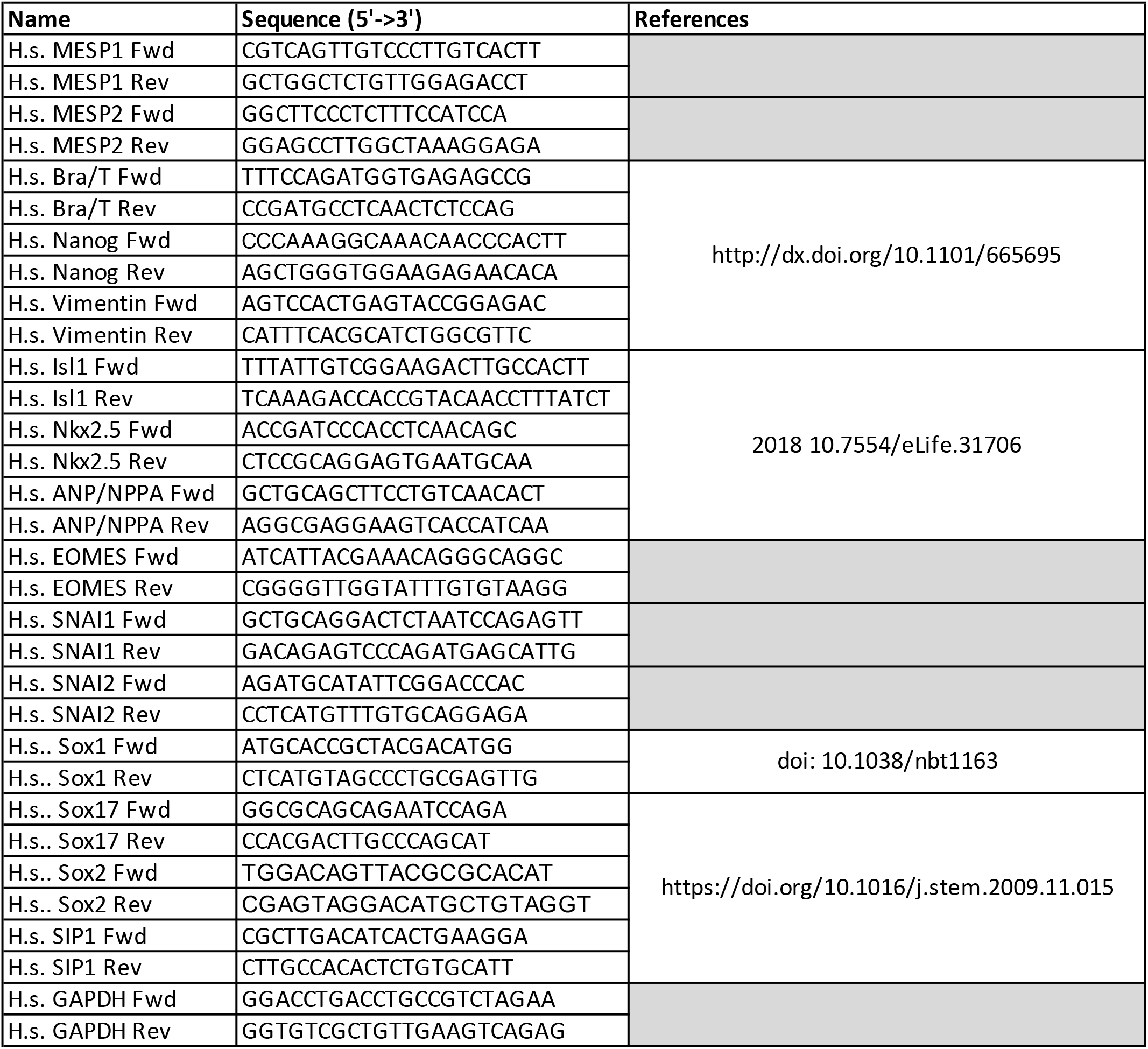
RT-qPCR primers

### Annexin V-APC assay

Protocol was adapted from the Annexin V apoptosis kit APC. Briefly, cells were collected as single cell suspension by incubation in gentle cell dissociation buffer for 8 min at 37°C. Pellet was washed in once in PBS and once in 1X binding buffer. Cells were resuspended in 100 μL 1X binding buffer and incubated 15 min at room temperature with 5 μL of Annexin-APC. Cells were washed in 1X binding buffer, resuspended in 200 μL of binding buffer and incubated with 5 μL of propidium iodide. Cells were passed through a 70 μm strainer prior to cytometry analysis using a 3-laser Fortessa flow cytometer.

### ATP release - Luciferase assay

hiPSCs were treated with CHIR-99021 as described previously plus DMSO or 10 μM Q-VD-OPH. Aliquots of culture medium (300 μl) were taken at indicated timepoints and mixed with 100 μL of 4X RealTime-Glo extracellular ATP assay reagent (Promega) reconstituted in RPMI 1640/B27(-Ins) medium. Technical triplicates of 100 μL were dispensed into a dark edged glass-bottom 96-well plate. Luminescence was measured after 30min using a HT-Synergy plate reader. Luminescence was subtracted for background.

A standard curve was obtained by serial dilution of ATP in RPMI 1640/B27(-Ins) media followed by the luciferase assay as described above. Simple linear regression was applied to transform luminescence values to ATP concentration.

### Statistical analysis

Datasets were analyzed using Prism8 (v.8.4.3) and tested for normality prior to applying the appropriate statistical test, as mentioned in each figure legend. Error bars represent S.D unless stated otherwise. Significance levels are given as follows: n.s. (not significant) : P > 0.05, *P ≤ 0.05, **P ≤ 0.01, ***P ≤ 0.001, ****P ≤ 0.0001.

All experiments were repeated at least three times independently as biological repeats unless stated otherwise.

Datasets are color-coded to reflect the variability between biological repeats.

## Acknowledgments

We thank Piyush Joshi for establishing the BAX/BAK DKO hiPSCs and Megan Rasmussen for the providing the SIM pictures of hiPSC-derived cardiomyocytes. We thank members of the Macara lab for discussion. This work was supported by GM070902 from NIGMS, CA197571 from the NCI (both to I.G.M) and by 1R35GM128915-01 (to V.G.).

## Contributions

Conceptualization, L.F., V.G. and I.G.M. ; Methodology, L.F., I.G.M. ; L.F. performed experiments and analyzed data. V.G. provided resources. L.F. prepared the figures. I.G.M and LF. wrote and edited the manuscript. I.G.M. supervised the work.

---

**Supp. Fig. 1.**
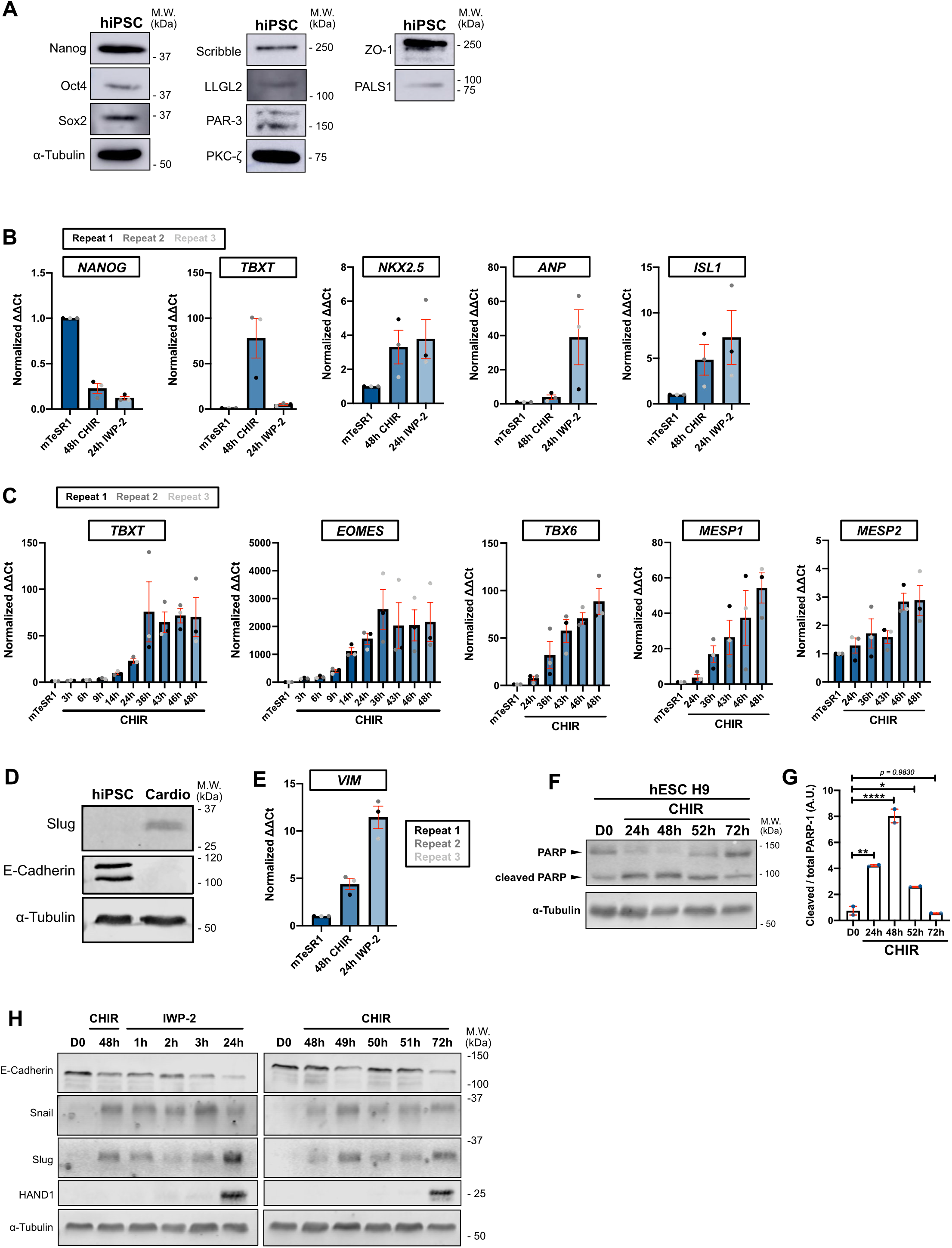
A)Immunoblot of WT hiPSC for pluripotency markers (Nanog, Oct4, Sox2), tight junctions marker (ZO-1), Crumb complex (PALS1), PAR complex (PAR-3, PKC-ζ), Scribble complex (LLGL2, Scribble) an α-Tubulin. Molecular weights (M.W.) are indicated in kDa. B) Relative gene expression of *NANOG, TBXT, NKX2.5, ANP, ISL1* before (mTeSR1) and during differentiation to mesoderm (CHIR) and cardiac mesoderm (IWP-2). ΔΔCt values were normalized to the mTseSR1 condition. Independent biological repeats are color-coded (n=3). Mean +/− S.E.M. C) Relative gene expression of *TBXT, EOMES, TBX6, MESP1, MESP2* before (mTesSR1) and during CHIR induction. ΔΔCt values were normalized to the mTseSR1 condition. Independent biological repeats are color-coded (n=3). Mean +/− S.E.M. D) Immunoblot of hiPSCs and hiPSC-derived cardiomyocytes obtained after applying the differentiation protocol described in Fig. 1B. Expression of EMT markers (Slug and E-Cadherin) were analyzed. Molecular weights (M.W.) are indicated in kDa. E) Relative gene expression of *VIM* was analyzed by qRT-PCR during cell conversion. ΔΔCt values were normalized to the mTseSR1 condition. Independent biological repeats are color-coded (n=3). Mean +/− S.E.M. F-G) Immunoblot of PARP cleavage in hESC H9 during CHIR treatment. Molecular weights (M.W.) are indicated in kDa (F). PARP cleavage was quantified by densitometry across 2 independent biological repeats (color-coded). Tukey’s multiple comparison was applied (* p ≤ 0.05, ** p ≤ 0.01, *** p ≤ 0.001 (G). H) Immunoblot comparing expression of EMT markers (E-Cadherin, Snail, Slug) and cardiac marker (HAND1) between the 2 protocols shown in Fig. 1A.

**Supp. 3.**
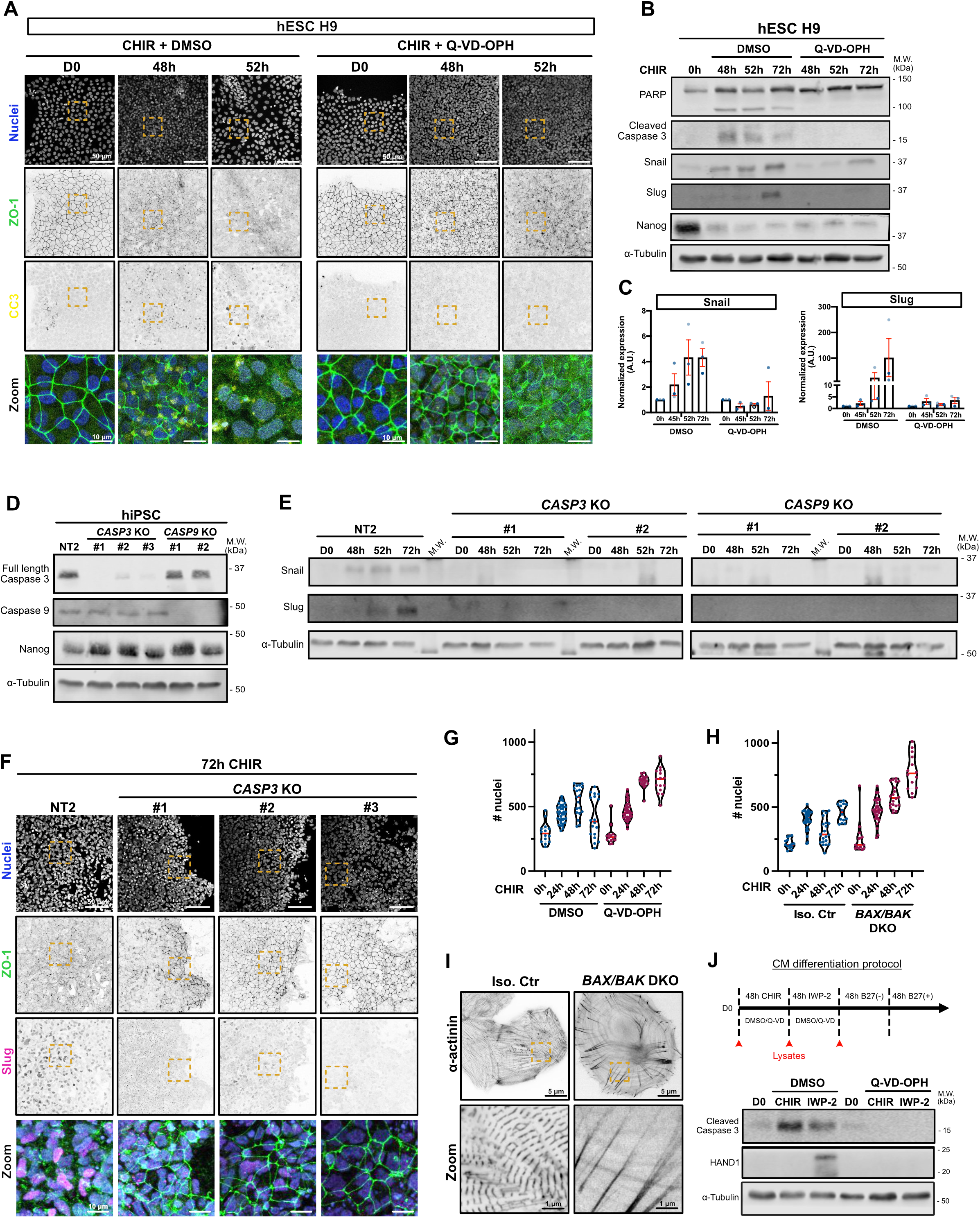
A) Representative immunofluorescence images of hESC H9 co-treated with CHIR and DMSO (left) or CHIR and 10 μM Q-VD-OPH (right) stained for ZO-1 (green), cleaved caspase 3 (yellow) and nuclei (blue). Maximum intensity projections are shown. Scale bar = 50 μm. Magnified area (yellow dotted square) is shown as a merge. Scale bar = 10 μm. B-C) Immunoblot analysis of hESC H9 co-treated with CHIR plus DMSO or 10 μM Q-VD-OPH. Molecular weights (M.W.) are indicated in kDa (B). Normalized expression of Snail and Slug was quantified by densitometry across 3 independent biological replicates (color-coded). Mean +/− S.E.M. (C). D) Immunoblot of *CASP3* and *CASP9* KO cell lines (non clonal). Knock-out validation was performed by probing for Caspase 3 and Caspase 9 expression, as well as Nanog as a stem cell marker. Molecular weights (M.W.) are indicated in kDa. E) Immunoblot of control Non Targeted (NT2) and *CASP3* and *CASP9* KO cell lines, analyzed for Snail and Slug expression upon CHIR induction. Molecular weights (M.W.) are indicated in kDa. F) Representative immunofluorescence images of Non targeted (NT2) and *CASP3* and *CASP9* KO hiPSCs, induced 72h with CHIR and stained for ZO-1 (green), Slug (magenta) and nuclei (blue). Maximum intensity projections are shown. Scale bar = 50 μm. Magnified area (yellow dotted square) is shown as a merge. Scale bar = 10 μm. G-H) Violin plots representing numbers of nuclei over time for WT hiPSCs co-treated with CHIR and Q-VD-OPH (G) or BAX/BAK DKO hiPSCs treated with CHIR. (Median: plain red line – Quartiles: black dotted lines). Three independent biological repeats (5-12 random fields of view/repeat). I) Immunofluorescent images of isogenic control and *BAX/BAK* DKO hiPSC-derived cardiomyocytes, plated at low density and stained for alpha-actinin. Scale bar = 5 μm. Magnified area (yellow dotted square) is shown. Scale bar = 1 μm. J) Immunoblot analysis of hiPSC treated with CHIR and IWP-2 co-treated or not with 10 μM Q-VD-OPH and probed for cleaved caspase 3 and cardiac marker HAND1. Cell lysates were collected as indicated on the timeline (red arrows).

**Supp. Fig. 4.**
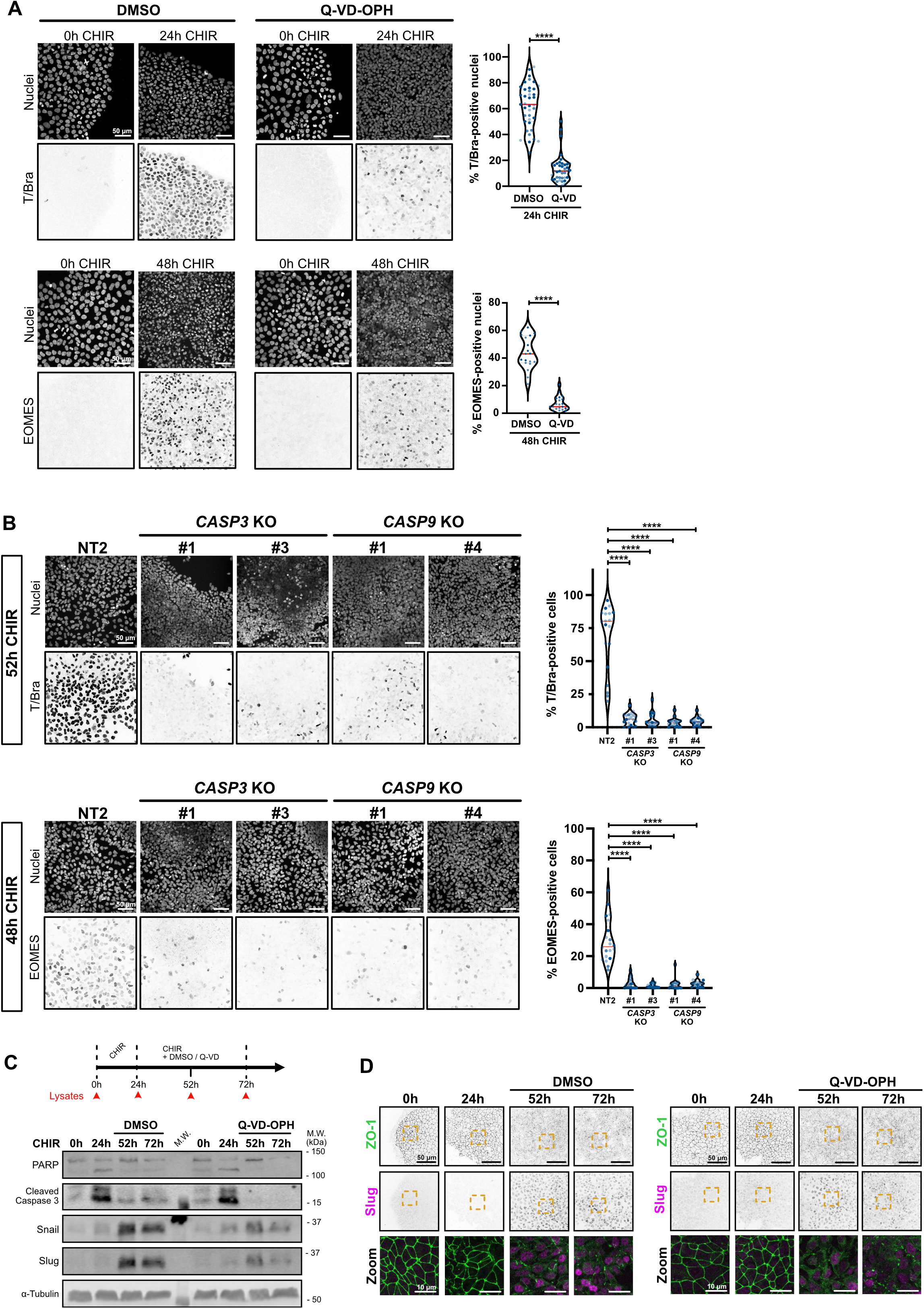
A) Representative immunofluorescence images of WT hiPSCs co-treated with CHIR +/− Q-VD-OPH and stained for T/Bra (top), EOMES (bottom) and nuclei, before (0h) and after CHIR treatment. Maximum intensity projections are shown. Scale bar = 50 μm. Violin plots summarize quantification of the percentage of T/Bra-positive (top) and EOMES-positive (bottom) nuclei after CHIR treatment (Median: plain red line – Quartiles: black dotted lines). Independent biological repeats are color-coded (n=3, 10-13 random fields of view/repeat for T/Bra and n=2, 10 random fields of view/repeat for EOMES). Mann-Whitney test was applied (**** p ≤ 0.001). B) Representative immunofluorescence images of control NT2, *CASP3* and *CASP9* KO hiPSCs stained for T/Bra (top), EOMES (bottom) and nuclei after CHIR treatment. Maximum intensity projections are shown. Scale bar = 50 μm. Violin plots summarize quantification of the percentage of T/Bra-positive (top) and EOMES-positive (bottom) nuclei after CHIR treatment (Median: plain red line – Quartiles: black dotted lines). Independent biological repeats are color-coded (n=3, 5-7 random fields of view/repeat. Kurskal-Wallis test was applied (**** p ≤ 0.001). C) Immunoblot analysis of hiPSCs treated with CHIR only for 24h, before adding CHIR +/− 10 μM Q-VD-OPH for another 28h and 48h with CHIR (respectively 52h and 72h timepoint). Treatment timing is indicated on the timeline and lysate collection is depicted by red arrows.). Molecular weights (M.W.) are indicated in kDa. D) Representative immunofluorescence images of hiPSCs treated with CHIR for 24h before addition of 10 μM Q-VD-OPH or DMSO. Cells were cultured for another 28h or 72h and stained for ZO-1 (green) and Slug (magenta). Maximum intensity projections are shown. Scale bar = 50 μm. Magnified area (yellow dotted square) is shown as a merge (bottom row). Scale bar = 10 μm.

**Supp. Fig. 5.**
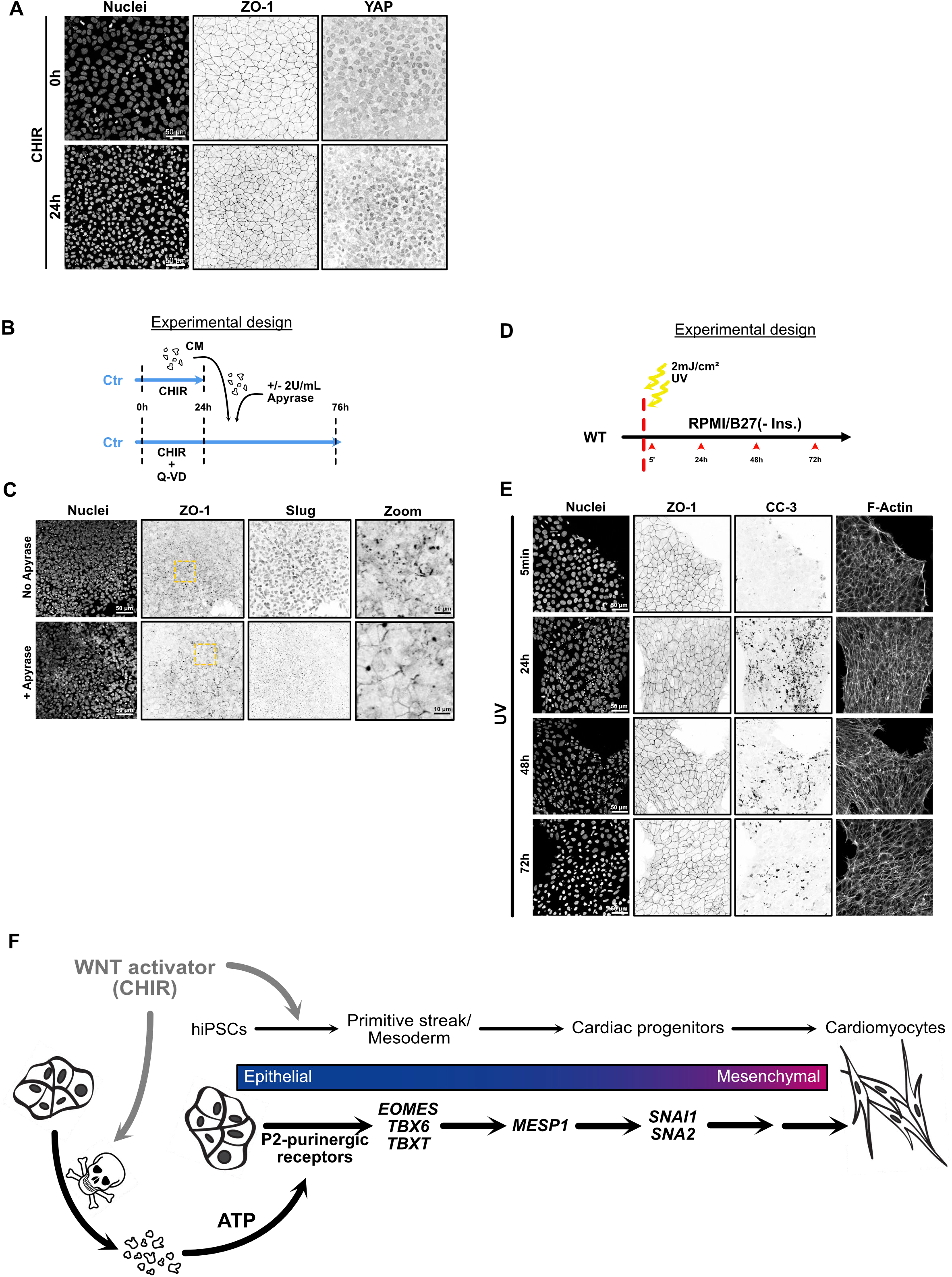
A)Representative immunofluorescence pictures of WT hiPSCs fixed after CHIR treatment and stained for ZO-1, YAP and nuclei. Maximum intensity projections are shown. Scale bar = 50 μm. B) Timeline of apyrase-treated conditioned media (CM) experiment. C) Representative immunofluorescence pictures of WT cells fixed after adding apyrase-treated CM as depicted in (B). Cells were stained for ZO-1, Slug and nuclei. Maximum intensity projections are shown. Scale bar = 50 μm. Magnified area (yellow dotted square) is shown for the ZO-1 channel. Scale bar = 10 μm. D) Timeline for UV exposure. WT hiPSCs were irradiated with 2mJ/cm2, kept in RPMI/B27(-Ins) media without CHIR and fixed at the indicated timepoint (red arrowheads). E) Representative immunofluorescence images from UV-irradiated hiPSC, stained for nuclei, ZO-1, cleaved caspase-3 (CC-3) and F-actin. Maximum intensity projections are shown. Scale bar = 50 μm. F) Working model

## Movie legends

**Supplementary movie 1**

Phase contrast timelapse of hiPSC-derived cardiomyocytes obtained using the GiWi differentiation protocol. Spontaneous beating was observed 12 days after protocol initiation and immature cardiomyocytes were maintained in RPMI/B27 (+ Ins.).

**Supplementary movie 2**

Timelapse imaging of mEGFP-TJP1 knock-in hiPSCs, starting 40h after CHIR-99021 treatment. Scale bar = 50 μm. Maximum intensity projections are shown.

**Supplementary movie 3**

Phase contrast timelapse of control (NT2) and *MESP1* knockout (#1 and #2) hiPSC-derived cardiomyocytes. Movies were recorded 12 days after GiWi protocol initiation using an EVOS FL microscope (Obj. x10). Scale bar = 400 μm.

**Supplementary movie 4**

Phase contrast timelapse of control (Ctr) and *BAX/BAK* DKO hiPSC-derived cardiomyocytes. Movies were recorded 12 days after GiWi protocol initiation using an EVOS FL microscope (Obj. x10). Scale bar = 400 μm.

